# Immunological contributions to age-dependent variations in behavioural responses to cutaneous inflammation

**DOI:** 10.1101/2022.07.15.500174

**Authors:** Emma Dayman, Andrew Bennett, Gareth J. Hathway

## Abstract

Systemic responses to immune challenge are immature at birth. However exposure to experimental inflammogens are able to produce an immunologic response which is characterised by swelling and oedema but, unlike in adults, does not result in sensory hypersensitivity. We sought to investigate whether the lack of nociceptive hypersensitivity was as a result of altered hemapoietic immune cell recruitment to the site of inflammation and/or differences in the cytokine and chemokine profile released by tissue invading cells. Postnatal (day of birth) and young adult (40-days old) Sprague-Dawley rats were used. Inflammation was induced by s.c. injection of Complete Freunds Adjuvant (CFA) unilaterally into the one hind paw. Mechanical withdrawal thresholds were measured before and after injection (2-168hrs). In adults a significant hyperalgesia was evoked which was absent in neonates. Immunohistochemical analysis of invading immune cells present in the perfusion fixed skin showed that although total cell numbers in the paw were the same in both age groups, neonates recruited more cells positive for both cell surface markers CD68 and Mannose-receptor (MR) whereas adults recruited significantly more cells positive for MR alone. There were no differences in neutrophil recruitment (as measured with H&E staining). TaqMan qPCR demonstrated that the temporal profile of cytokine production in the skin differed between ages with neonates responding faster than adults and that neonates produced significantly more IL-1b and IL-27 then adults who expressed significantly more IL-6 and IL-10. This study illustrates that in neonates the cell recruitment and cytokine profiles are markedly different to those seen in adults; this may in part explain why behavioural responses to inflammation are suppressed relative to adults.

## 1. Introduction

Pain impacts the lives of people across the life-course, yet the way in which pain is detected and processed is different a different life stages. The perception of pain is especially hard to assess in non-verbal populations, including the very young and older people with disabilities. It is therefore important to understand how age-related alterations in biological processes such as tissue responses to damage or infection alter the way in which pain is detected, processed and perceived.

Nociceptive pathways are subject to major reorganisation and refinement during early life(Fitzgerald, 2005). Noxious stimuli evoke uncoordinated nocifensive responses in young animals (<21days old in rodents) coupled with significantly lowered mechanical and thermal withdrawal responses when compared with adults (Falcon et al., 1996; FitzgeraId et al., 1988; Fitzgerald and Jennings, 1999). Concurrently neonates have an altered response to immunological challenge. The interactions between neuronal and immunological systems and the impact these have on altered pain sensitivity has started to gain increased attention in recent years (Zouikr et al., 2016; Zouikr and Karshikoff, 2017). Moreover, substantial evidence indicates that tissue damage and accompanying pain during the neonatal period profoundly affects the maturation of pain pathways, with a lasting impact on sensory thresholds and pain perception(Fitzgerald and Walker, 2009; Walker et al., 2016). Despite the clear overlap in these physiological processes, the role of the systemic immune system in pain mechanisms and how immunological responses are different in early life has not been explored thoroughly.

The neonatal immune system differs significantly from the mature immune system. It is considered somewhat immunodeficient, with many components functioning differently to the adult during the transition from the protective intra-uterine environment to the outside world (Levy, 2007). In 1996, three key papers were published that demonstrated that the adaptive immune system of neonates is competent and under certain conditions is able to mount T-cell responses comparable to levels seen in adults (Forsthuber et al., 1996; Ridge et al., 1996; Sarzotti et al., 1996). However, the innate immune system at birth and in the perinatal period is adapting from existing in an immunologically protected environment in utero in which anti-inflammatory responses predominate to prevent inflammatory responses to maternal antigens. This Th2 predominating, anti-inflammatory phenotype leads to a distinct immunological response, immunotolerance and low control over infections (Yu et al., 2018). Neutrophil numbers are lower in the neonate and those that are present are of subdued quality and function (Yu et al., 2018). They express lower levels of Pathogen Activated Molecular Patterns (PAMPs) such as TLR2 and TLR4. Neonatal antigen presenting cells (APCs) such as macrophages and dendritic cells (collectively monocytes) show lower levels of expression of MHCII, CD80/86 and TLRs than adults.

In adult rats, an inflammatory immune response can be initiated via a subcutaneous injection of an experimental inflammogen such as complete Freund’s adjuvant (CFA). This induces a local immune response via the activation of toll-like receptors (TLR-2, −4 and −9), as well as IL-1R on tissue resident immune cells (Coffman et al., 2010). Initiation of this cascade provokes the infiltration of neutrophils, macrophages and the activation of mast cells at the site of injection, which release various inflammatory mediators including histamine, pro-inflammatory cytokines (e.g. TNFα, IL-1β, IL-6), vasodilators such as bradykinin and prostaglandins including PGE_2_. This generates typical symptoms of oedema, redness and pain. In neonates, this acute oedema and redness also occur in response to inflammogens and persists for approximately 1-2 weeks, however, the sensory hypersensitivity seen in adults does not develop (Walker et al., 2003).

Here we investigated whether differences in the innate immune response between adults and neonates could account for differences in pain behavioural responses to inflammation.

## 2. Materials and methods

### 2.1 Animals

All experimental procedures in animals were licenced and conducted in accordance with the UK Home Office Animals (Scientific Procedures) Act 1986 (ASPA 1986) and the International Association for the Study of Pain (IASP) and ARRIVE guidelines (Percie du Sert et al., 2020). Studies were approved by the ethical review board within the University of Nottingham and research was carried out under PPL PB3DA999F and PIL I860D2962.

Experiments were carried out on rats aged postnatal day (P) 1-2 and ‘adult’ (>P40) Sprague Dawley rats that were purchased from Charles River, UK. Pups were housed with their dams in individually ventilated cages in an in-house animal facility. Free access to food and water was available throughout. Adult animals were group housed (4 per cage) in conventional wire top cages in with free access to food and water throughout. All animals were kept on a 12:12h light dark cycle, with all studies taking place during the light cycle. Prior to beginning a study, the animals were habituated to the experimental equipment and the experimenter was blinded to the treatment of each group to ensure no bias throughout.

### 2.2 Anaesthesia

Isoflurane was chosen as the general inhalant anaesthesia (Abbott Laboratories Ltd, UK) as it provides an accurate and continuous depth of anaesthesia, which is easily adjusted if necessary during procedures. Animals were placed in a clear Perspex induction box and 3% isoflurane anaesthesia was delivered via an anaesthetic vaporiser (Burtons, Kent, UK), in a nitrous oxide and oxygen gas (BOC gases, UK) mixture in a ratio of 33:67. Once areflexic (confirmed by the pedal reflex), animals were transferred to a heat pad (37°C) in the prone position and a small animal nose cone was fitted allowing the delivery of isoflurane at 2.5%. Following recovery from anaesthesia, pups were then immediately transferred back to their home cage and dam to minimise stress caused by maternal separation whilst adult rats were returned to their homecage.

### 2.3 Induction of inflammatory models

#### 2.3.1 Carrageenan induced inflammatory pain model

λ-Carrageenan (Sigma-Aldrich) was immersed in 0.9% saline (NaCl) with a concentration of 2% carrageenan was used (Zhang and Ren, 2011).

#### 2.3.2 Intraplantar Administration of Substances

Once animals were anaesthetised and areflexia was confirmed so that no pain would be felt upon insertion of the needle, the animals paws were positioned so the plantar surface faced upwards. The plantar surface of the left hindpaw was disinfected with chlorohexidine. A 25-gauge; 16-mm needle (Becton, Dickinson and Company Ltd, Ireland) attached to a 1 ml hypodermic syringe (Terumo, New Jersey, U.S.A) was used to subcutaneously inject substances into the plantar skin of the paw. Once the procedure was completed, animals were allowed to recover in a separate cage and then transferred to its home cage.

### 2.4 Paw Withdrawal Threshold

The development of mechanical hyperalgesia in animals was determined by assessing changes in mechanical hindpaw withdrawal thresholds. Adults rats were placed in a clear Perspex box that had a fine stainless steel mesh floor, this was raised from the work surface allowing access from below. Calibrated von Frey monofilaments (Semmes-Weinstein monofilaments of bending forces 0.6, 1, 1.4, 2, 4, 6, 8, 10, 15 and 26g) were applied to the plantar surface of the hindpaw for three seconds, in ascending order of bending force until a withdrawal reflex was observed. Each paw was tested a total of three times and the average was calculated. All animals underwent two consecutive days of habituation to the testing procedure prior to baseline behavioural measurements. Neonates were gently restrained with one hand and monofilaments applied to the dorsal surface of hindpaws using the same method as in adult rats (Hathway et al., 2012). However, neonates did not undergo habituation in order to reduce the impact of stress due to maternal separation.

### 2.5 Haematoxylin and Eosin (H&E) staining

Skin sections were mounted onto gelatinised glass slides and sections were briefly washed in distilled water for 1 minute, followed by staining in Harris haematoxylin solution for ~5 minutes. After washing and dehydration H&E stained sections were observed with a Leica DM4000B upright light microscope, capable of imaging using a QImaging Micropublisher 3.3 RTV camera and Micro-Manager 1.4 software. Sections were imaged using a 50x oil / 0.9NA N-Plan and 100x oil /1.25 NA N-Plan. Fiji (Schindelin *et al*., 2012) was used to perform image analysis offline, including, adjustment of image brightness/contrast and cells counts. Images were processed identically and the same contrast enhancement was applied to each image.

### 2.6 Immunohistochemistry

#### 2.6.1 Tissue collection

Fresh and perfusion fixed skin sections were used in this study. For fresh tissue neonatal and adult rats were overdosed with an i.p. injection of sodium pentobarbital and death was confirmed by cervical dislocation. In adults, hindpaw skin was freshly collected from both the ipsilateral and contralateral paw and in neonates the entire paw was taken as the skin was too delicate to dissect. Tissue was stored at −80°C prior to processing. For fixed tissue both adult and neonatal rats were overdosed with an i.p. injection of sodium pentobarbital, then animals were transcardially perfused with ice cold PBS followed by 4% paraformaldehyde (Sigma). Skin was collected, as above, and then post-fixed overnight in paraformaldehyde at room temperature. Sections were then transferred to 30% sucrose and 0.01% azide solution for cryoprotection and stored at 4°C until processed. Hindpaw skin samples were sectioned on the freezing microtome (Leica, SM2010R) set at a nominal thickness of 30μm. Sections were collected onto 24 well plates filled with sucrose azide solution and stored at 4°C.

#### 2.6.2 Immunofluorescent staining

Skin sections were mounted onto gelatinised glass microscope slides and allowed to dry overnight at room temperature. Sections were washed three times for 10 minutes with 0.1M PBS. An antigen retrieval step was then incorporated into the protocol by washing the sections three times for 10 minutes with 10mM citrate buffer heated to 50°C. Sections were then washed a further three times for 10 minutes with 0.1M PBS. Sections were blocked with 6 % (v/v) goat serum in 0.1 M PBS/0.01% (v/v) Triton for 2 hours at room temperature followed by incubation with primary antibodies (Mouse monoclonal anti-rat CD68 (ED1) BIO-RAD, (1:100); rabbit polyclonal anti-rat Mannose Receptor Abcam (1:300)) overnight at room temperature. Both primary and secondary antibody dilutions were prepared in TTBS (Tris-Buffered Saline and Tween 20). Following primary antibody incubation, sections were washed then incubated with secondary antibodies (Goat anti-Mouse Alexa Fluor 488, ThermoFisher; Goat anti-Rabbit Alexa Fluor 568, ThermoFisher (both 1:500)) for 2 hours in a dark box at room temperature. Following incubation, sections were washed and then stained with 2μg/ml DAPI (4’,6-diamidino-2-phenylindole) in 0.1M PBS to stain cell nuclei. The sections were allowed to dry overnight at room temperature and were mounted in a Fluromount Aqueos (Sigma-Aldrich) mounting media prior to fluorescent microscopic visual analysis and quantification.

#### 2.6.3 Microscopy and image analysis

Immunofluorescent sections were observed with a Zeiss 200M inverted fluorescence microscope. Images were captured using Micro-Manager 1.4 software to control a CoolSNAP MYO with exposure time ranging from 0.5-2.0 seconds. Sections were observed using a 40x oil immersion lens (numerical aperture of 1.3). Fiji (Schindelin *et al*., 2012) was used to perform image analysis offline, including, adjustment of image brightness/contrast, cells counts and fluorescence. Images were processed identically and the same contrast enhancement was applied to each image. Four skin samples were imaged per animal (eight animals in total per group) with four fields of view per sample.

### 2.7 Taqman real-time polymerase chain reaction (RT-PCR)

#### 2.7.1 RNA extraction

For extraction of RNA from fresh paw skin, the Direct-zolTM RNA Miniprep Plus kit (Zymo Research) was useds as per the manufacturer’s instructions; 50mg of paw tissue was processed in 600μl of TRI Reagent.

#### 2.7.2 cDNA synthesis

Reverse transcription of RNA was performed using the AffinityScript QPCR cDNA Synthesis Kit (Agilent Technologies). Total RNA e (250ng) was rverse transcribed according to the manufacturers instructions. Samples were stored at −20°C prior to future use.

#### 2.7.4 RT-PCR

#### 2.7.5 Quantification of target gene expression

The relative standard curve method based on Taqman qRT-PCR was used for quantifying gene expression. The slope of the standard Assays were deemed acceptable if there was <0.5 cycle threshold (Ct) difference between the values of a triplicate. Slopes within the range −3.2 to −3.6 and r^2^ between 0.95-0.99 were deemed acceptable. A standard curve was included for each plate run and each gene expression studied. Target gene expression was quantified relative to the expression of the reference gene, GAPDH.

#### 2.7.7 Primers and probes

Primers and probes for gene expression studies were designed using primer express 3 software (Life Technologies),probes and primers were synthesised by Eurofins Scientific. Primer and probe sequences are shown in Table 1.

**Table 1:**
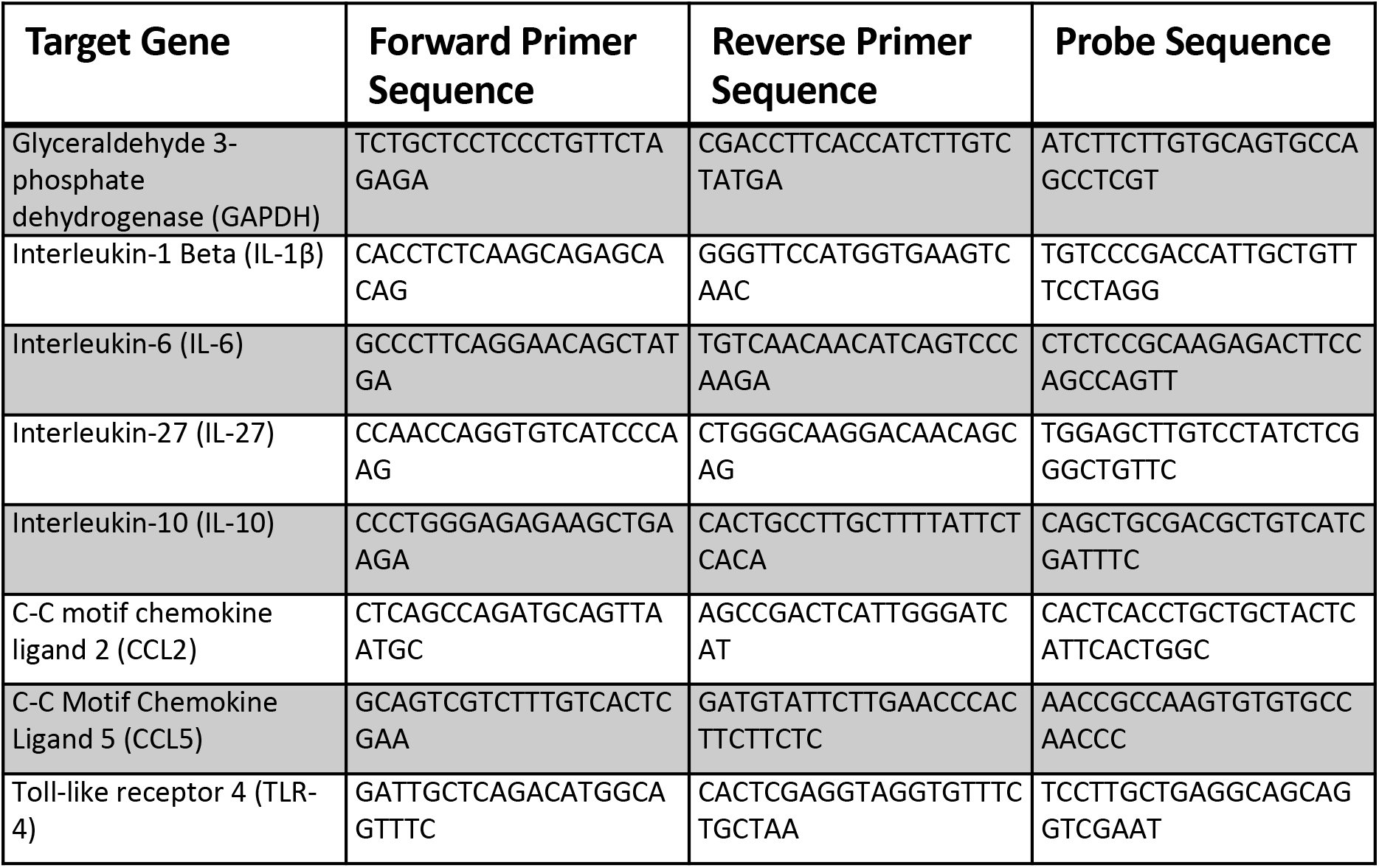
Sequences of primers and probes used in TaqMan PCR.

### 2.8 Statistical analyses

Data are represented as mean ± standard error of mean (SEM). Statistical significance was determined using appropriate tests (see below) with significant differences being designated when P<0.05. All data were analysed for normality of distribution using a Shapiro-Wilk normality test. Changes in PWTs were compared using 2-way repeated measures ANOVA with Bonferroni’s multiple comparison test. Analyses compared changes in pain behaviour over time with changes in the control group. Analysis of immunohistochemistry quantification was conducted using one-way ANOVA with Bonferroni’s multiple comparison test, where n = 4 per time point for each age group. PCR data was analysed using one-way ANOVA with Bonferroni’s multiple comparison test, where n = 6 per time point for each age group.

## 3. Results

### 3.1 Neonatal cutaneous inflammation does not induce changes in paw withdrawal threshold when compared with adult nociceptive testing

Cutaneous inflammation was induced via the intraplantar administration of CFA in neonatal (P1) and young adult (>P40) rats. In both groups this produced substantial swelling and erythema of the entire hindpaw, extending to the ankle, which resolved within 7 days and was not present in control groups. In young adult rats inflammation was accompanied by significant reductions in PWT from 2 hours post-injection only in the ipsilateral paw (F (9, 380) = 14.26, P<0.0001) (Figure 1b). However, neonates, did not show any accompanying alterations in mechanical PWT (Figure 1a). We also sought to establish the magnitude of swelling that accompanied CFA induced inflammation in both age groups. In both neonates and adults CFA injection was followed by swelling of the foot, which did not extend beyond the ankle in both ages. Quantifying this was problematic as plethysmometry in neonates could not be achieved and other methods such as measuring paw circumference with surgical suture and a ruler were highly variable. We therefore performed H&E staining (Fig 1c) and measured the number of cells which were present in skin sections of both ages. In both ages there were significant increases in cell counts at 2 and 6 hours post-CFA compared to controls (Figs 1d&e) however this did not differ between the ages. These data show that whilst behavioural hypersensitivity does not develop in neonates following inflammation, immune cell invasion in response to CFA was similar. Behavioural differences therefore are likely to be the result of more subtle changes in the immune response to experimental inflammogens.

**Figure 1:**
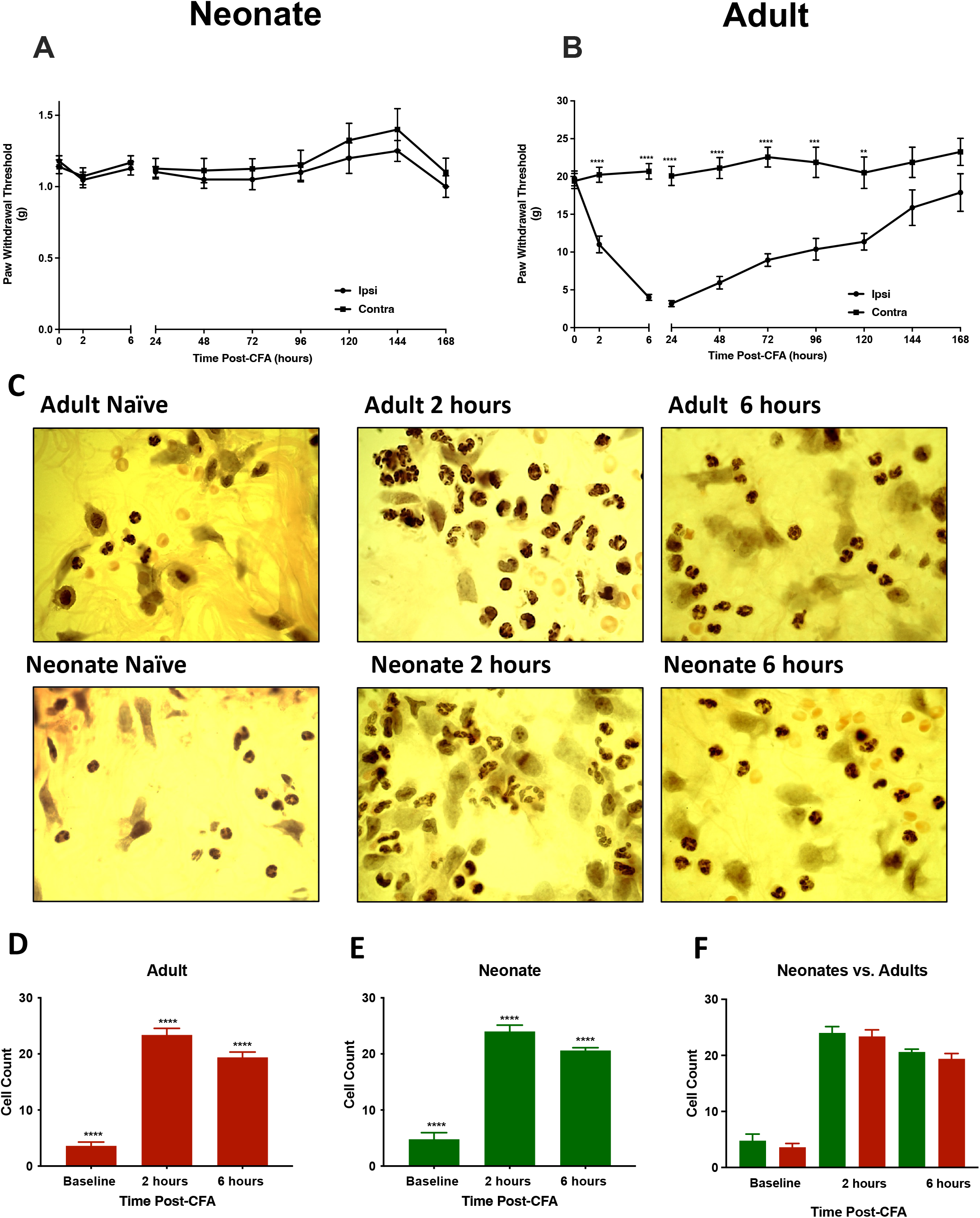
Subcutaneous CFA evokes different outcomes on mechanical withdrawal thresholds in adult and neonatal rats. In neonatal rats (A) CFA fails to evoke any behavioural sensitivity at any of the timepoints. However in adult rats (B) the inflammogen evokes a signficant hyperalgesia when compared to the contralateral paw. This is evident from 2hours until 120hours post-inection. Data displayed are mean ± S.E.M. Comparisons between ipsilateral and contralateral paws were made using a Two-way Repeated Measures ANOVA with Dunnett’s Multiple Comparison test. **P<0.01, ***P<0.001, ****P<0.0001. H&E staining of skin sections in response to CFA (C). The number of cells in the paw increased in both adult (D) and neonatal (E) animals compared to baseline controls but did not differ between the ages (F). Data displayed are mean ± S.E.M. ****P<0.0001 One-way ANOVA with Dunnett’s Multiple Comparison test.

### 3.2 Phenotype of infiltrating Immune cells differs between neonates and adults in response to cutaneous inflammation

We hypothesised that one potential explanation for the lack of behavioural hypersensitivity following inflammatory challenge in neonates was a difference in the level of M2 macrophages provoked to invade the injured tissue. We assessed this using anti-bodies directed against the mannose receptor (MR) a marker of M2 cells as well as CD68 a general marker of macrophages.

#### 3.2.1 The response of adult animals

Adult samples showed significant increases in the total number of immune cells at the site of inflammation at every time point post-CFA injection (Fig 2a and Fig 3) (F (5, 313) = 175.6, P<0.0001), with the biggest number of immune cells present at the 24 hour timepoint. CD68^+^MR^+^ cells were significantly increased from the 24hr timepoint compared to control animals (Fig 2d) (F (5, 313) = 38.6, P<0.0001). CD68 positive, MR negative cells remained low at 2 and 6 hours post-CFA; they increased significantly at 24 hours. One week after CFA, the mean number of CD68^+^ cells were significantly elevated compared with controls (Fig 2b) (F (5, 313) = 19.83, P<0.0001). Cells expressing only MR did not differ at 2 or 6 hours compared with control animals. From 24 hours through to 1 week, MR^+^ cell numbers were significantly increased (Fig 2c) (F (5, 313) = 51.05, P<0.0001).

**Figure 2:**
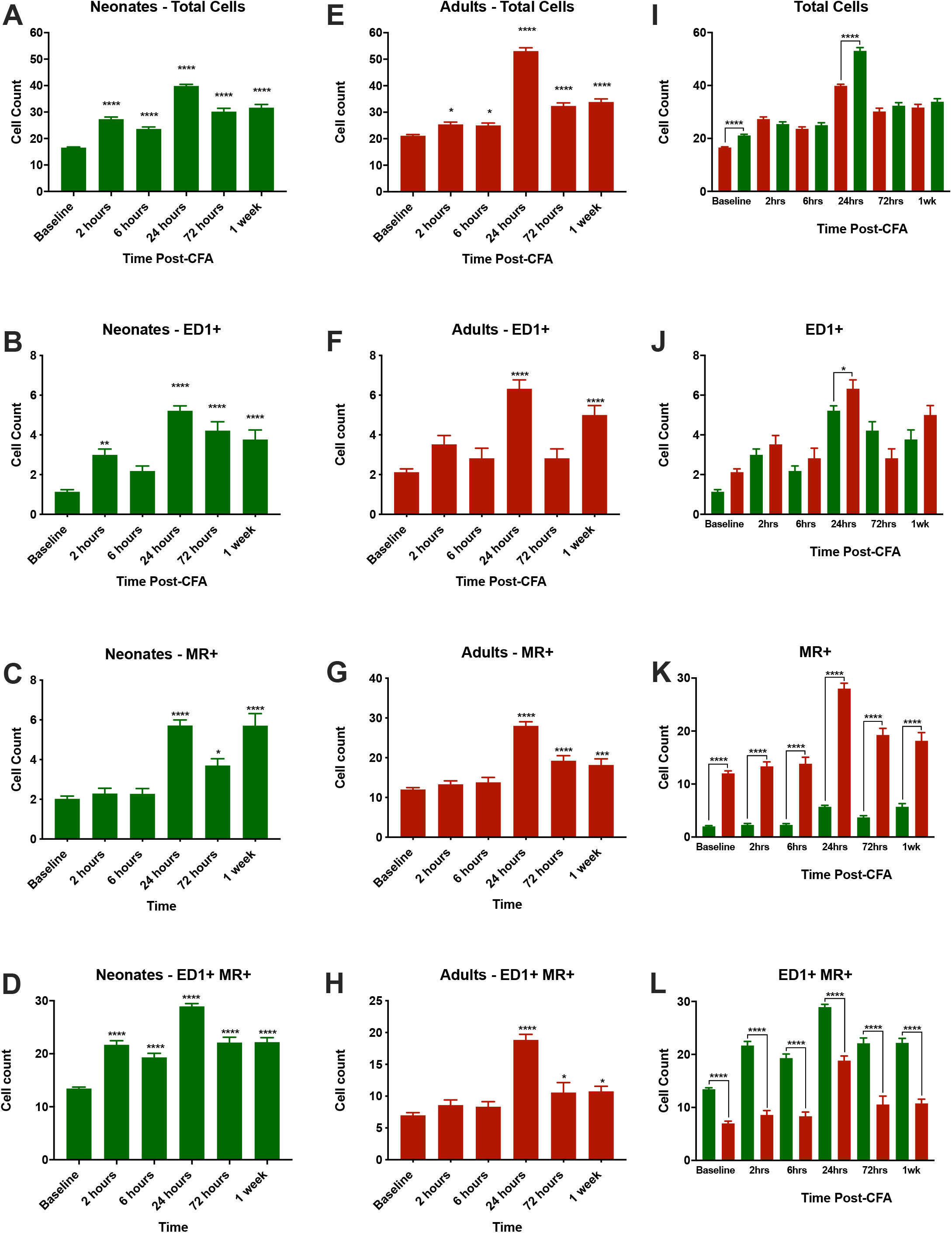
Immune cell infiltration of CFA exposed skin changes with age. Experimental inflammation induced by s.c. injection of CFA resulted in subtly different recruitment of monocytic cells. The total number of cells present in the skin significantly increased in both neonates (A) and adults (E) but there were limited differences between the age groups (I). CD68 positive monocytes were significantly increased in neonates from 2 hours post-CFA (B) but only increased from 24 hours in the adult (F). At the 24gr timepoint there were significantly more CD68 positive cells in adults than in neonates (J). The number of MR positive cells increased in both neonatal (C) and adult (G) from 24 hours compared to baseline, however there were always significantly more of these cells in adults compared to neonates at all timepoints (K). However cells which were positive for both CD68 and MR were significantly increased at all timepoints post-CFA in neonates (D) but at only 24 hour, 72 hour and 1 week post-CFA in adults (H). This cell type were far more common in the neonatal skin compared to the adult skin at all timepoints (L). Comparisons between timepoints within an age group and between age-groups at a given timepoint were made with One-way ANOVA with Bonferroni Posttest. *P<0.05, **P<0.01, ***P<0.001, ****P<0.0001.

**Figure 3:**
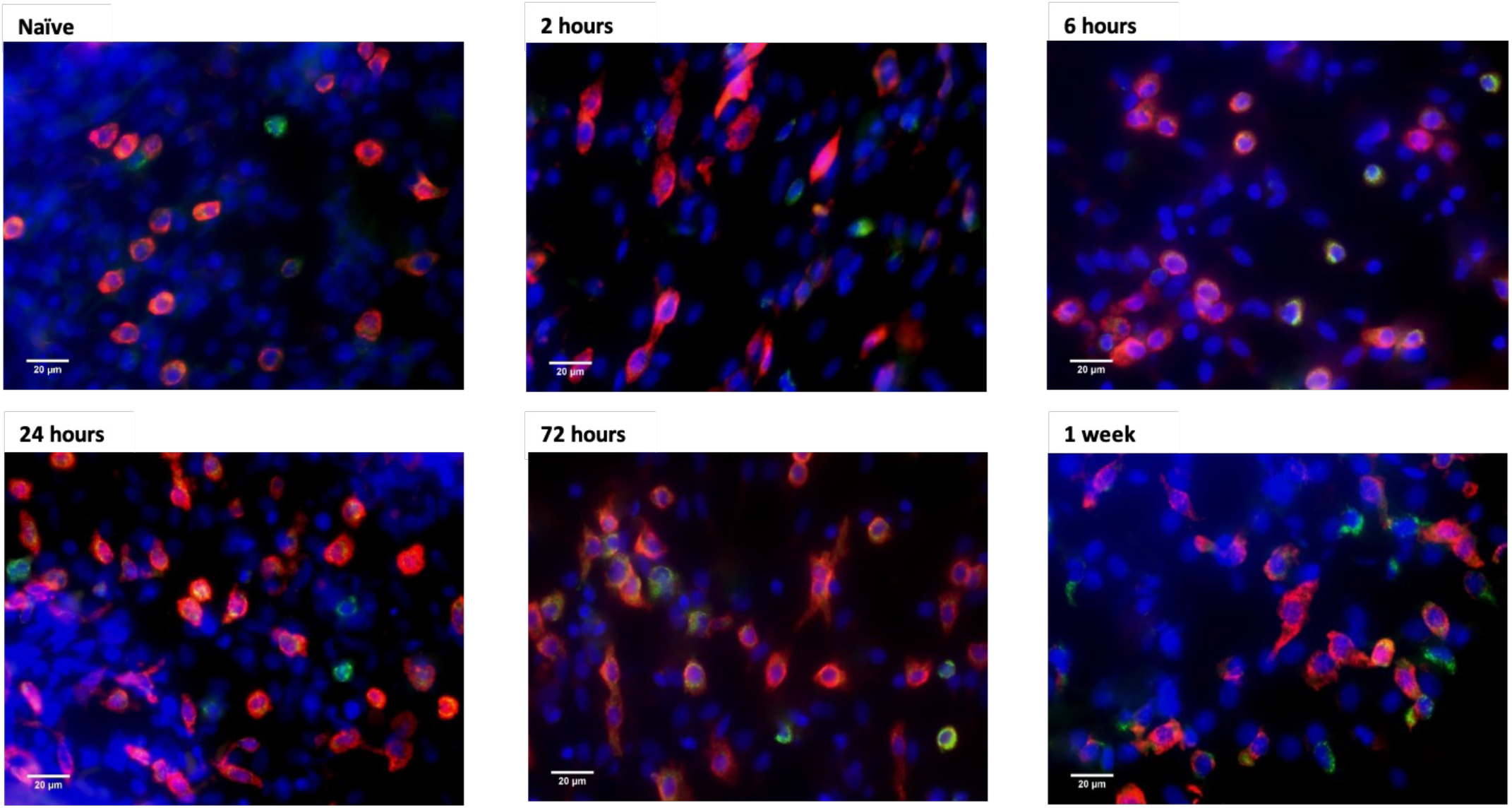
Neonatal immune cells undergo a change in shape following CFA injection. Immunohistochemical staining of CD68 (green), MR (red) and DAPI (blue) was used to assess morphological changes in immune cells in neonatal skin following s.c. CFA injection. Prior to CFA injection cells appear to have a rounded, regular appearance which changes to a more irregular shape, with processes as the period after CFA injection increases. To 24 hours. At the 72 hour timepoint cells appear to return to a morphology more consistent with the pre-CFA state and this process in complete by 1 week. Images were observed a 40X magnification, white bars indicate 20um.

#### 3.2.1 The response of neonatal animals

Within the neonatal group, there were significant increases in the total number cells at the site of injection at all time points compared with control animals (Fig 2e and Fig 4) (F (5, 690) = 199.5, P<0.0001). Within this population of infiltrating cells the number of CD68^+^MR^+^ macrophages significantly increased across all time points compared to control animals (Fig 2h) (F (5, 690) = 107.2, P<0.0001). Cells which only expressed CD68 were significantly increased at 2, 24, 72 hours and 1 week timepoints (F (5, 690) = 36.49, P<0.0001)(Fig 2f). Cells which were only MR^+^ significantly increased from 24 hours through to 1 week (F (5, 690) = 29.18, P<0.0001)(Fig 2g).

**Figure 4:**
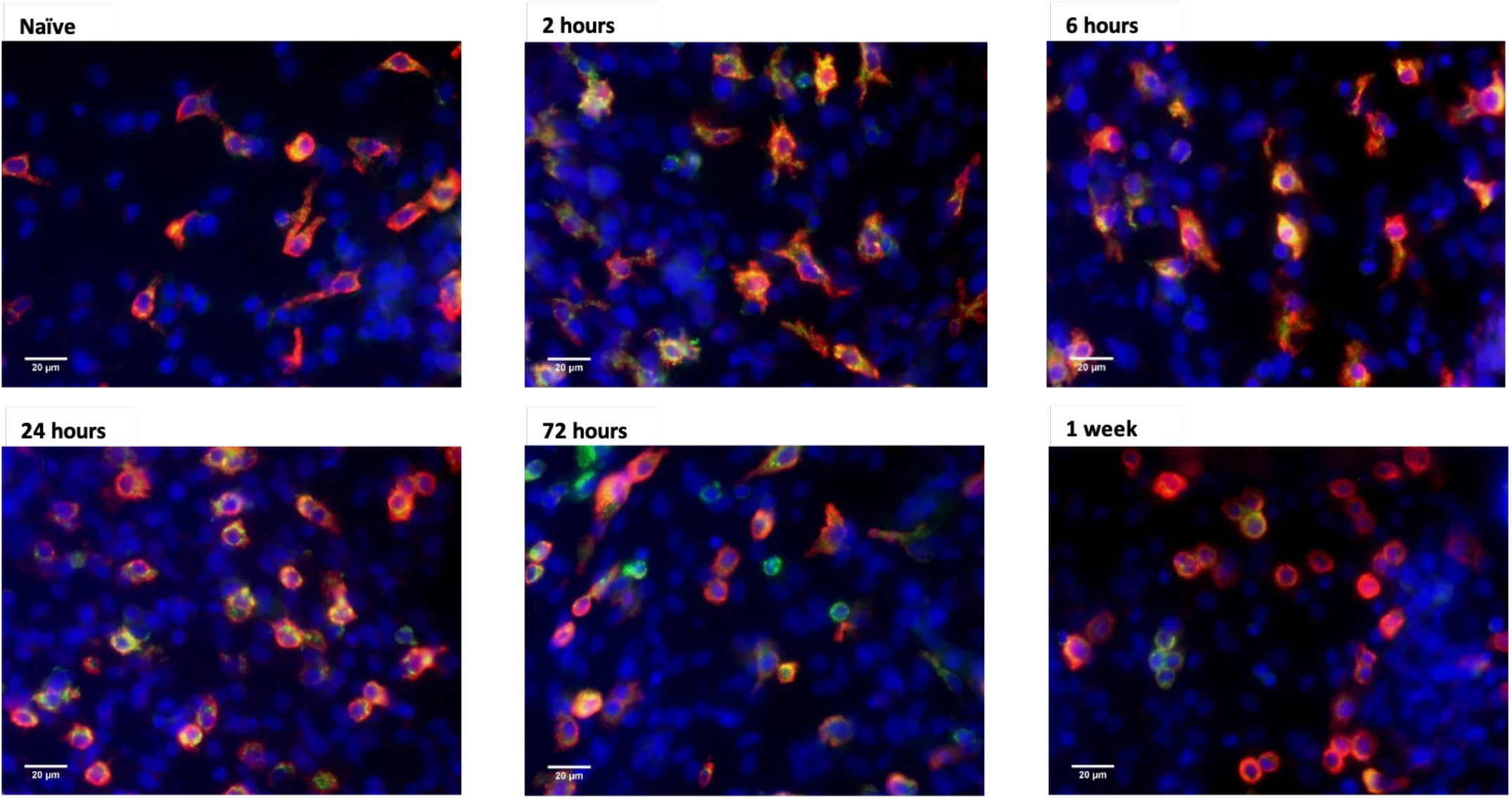
Adult immune cell morphology is unchanged following CFA injection. Assessment of adult immune cell morphology in adult skin samples post-CFA injection were made using immunohistochemistry. CD68 (green), MR (red) and DAPI (blue) show that cell morphology remains consistent throughout the 1 week period post-CFA injection. Images were observed a 40X magnification, white bars indicate 20um.

#### 3.2.3 Neonatal immune cells undergo a change in phenotype whilst adult cells do not

The phenotype/morphology of invading cells might be indicative of a change in function or expression profiles which impacts on behavioural responses to inflammogens. Immunofluorescence microscopy highlighted possible morphological differences in the immune cells between the neonatal and adult immune cells (Figs 3 and 4). This might be indicative in a difference in function. Overall, adult immune cells appeared regular and circular throughout (Fig 3). The shape of the cells over the time course of the adults remained fairly consistent. In neonates (Fig 4) cells in the naïve neonate group express high levels of MR and appear more circular and regular. At 2 and 6 hours post-CFA injection, there was an increased number of overall cells at the site of inflammation. Cells underwent a distinct change in shape, they were less regular, with increased co-expression of CD68 and MR expression. At 24 hours, cells began to return to their original, pre-CFA appearance. By 72 hours to 1 week there are fewer cells at the site of inflammation and these cells are returning to their usual regular/circular shape.

#### 3.2.4 Age significantly impacts on invading immune cell phenotype post-CFA inflammation

Comparisons between the two age groups highlighted clear differences in CD68 and MR expression over the timecourse of the post-CFA period. In control animals there are significantly more tissue residing monocytes/macrophages in neonates compared with adults (Fig 2j). The only time point that showed significant differences in the total number of immune cells at the site of inflammation was 24 hours (Fig 2k) (F (11, 1003) = 170.4, P<0.0001). Neonates have increased expression of CD68 MR^+^ cells at all time points compared with adults (Fig 2l) (F (11, 1003) = 121.1, P<0.0001). CD68 expression was largely not significantly different between control neonates and adults, the only time point where significant differences were seen for CD68 expression was 24 hours, (Fig 2m) (F (11, 1003) = 25.57, P<0.0001). Clear differences in the numbers of MR^+^ cells were observed. Adults had significantly higher numbers of cells expressing of MR at all time points compared to neonates, including higher expression of MR^+^ cells in control animals (Fig 2n) (F (11, 1003) = 199.5, P<0.0001). These data show that the response to cutaneous inflammation is different in neonates and adults. Whilst total number of invading cells is comparable between age groups, adult cells express more MR whereas neonatal cells co-express both MR and CD68. This difference is cell surface markers indicates differences in gene-expression profiles at these different ages.

### 3.4 Gene expression differences in cytokine and chemokine expression at the site of inflammation in neonates and adults

Haemopoietic cells invading tissue following injury secrete a range of neuroactive substances such as cytokines and chemokines which sensitise sensory fibres. We hypothesised that the absence of behavioural sensitisation following CFA administration in neonates may be the result of differences in gene expression between the age-groups and therefore secretion of these neuroactive substances. RT-PCR was used to study gene expression of the pro-inflammatory cytokines IL-1β and IL-6 as well as the anti-inflammatory cytokines IL-10 and IL-27 which have previously been shown impact nociception in adults. We also used the same approach to measure the chemokines CCL2 and CCL5. Tissue was freshly collected from neonatal and adult rats 2 hours, 6 hours, 24 hours, 72 hours and 1 week post-CFA injection to study gene expression post-CFA injection.

#### 3.4.1 CFA induced changes in cytokine gene expression differs between adults and neonates

In adult animals, the expression levels of pro-inflammatory cytokines IL-1β and IL-6 followed a similar pattern. IL-1β expression was increased by 6 hours post-CFA injection, while this was significant (P<0.05) there was significantly higher expression levels of IL-1β at 24 hours (P<0.0001) (Control: 0.17 ± 0.02, 6 hours: 0.41 ± 0.11, 24 hours: 2.25 ± 0.07 F (5, 27) = 140.7) (Fig 5a). By 72 hours expression levels were comparable to those in control animals. IL-6 expression was significantly increased at 6 hours and 24 hours post-CFA injection (Control: 0.09 ± 0.01, 6 hours: 4.02 ± 0.67, 24 hours: 2.61 ± 0.23 F (5, 29) = 29.77, P<0.0001) (Fig 5b). By 72 hours expression levels had reduced to a level similar to those in Control animals. IL-10 expression was only significantly increased at 24 hours in adult tissue. There were no changes to the expression of IL-10 at any other time point post-CFA injection (Control: 0.57 ± 0.06, 24 hours: 4.20 ± 0.15. F (5, 27) = 290.4, P<0.0001) (Fig 7c). IL-27 expression in adult animals is different to that of neonates. In adults, expression of IL-27 was supressed significantly at 2 hours and 6 hours (Control: 1.4 ± 0.39, 2 hours: 0.2 ± 0.02, 6 hours: 0.3 ± 0.09. F (5, 29) = 16.01, P<0.0001). At 24 hours there was a significant increase in IL-27 expression (24 hours: 2.48 ± 0.33 F(5,29)=16.01 P<0.01).

**Figure 5:**
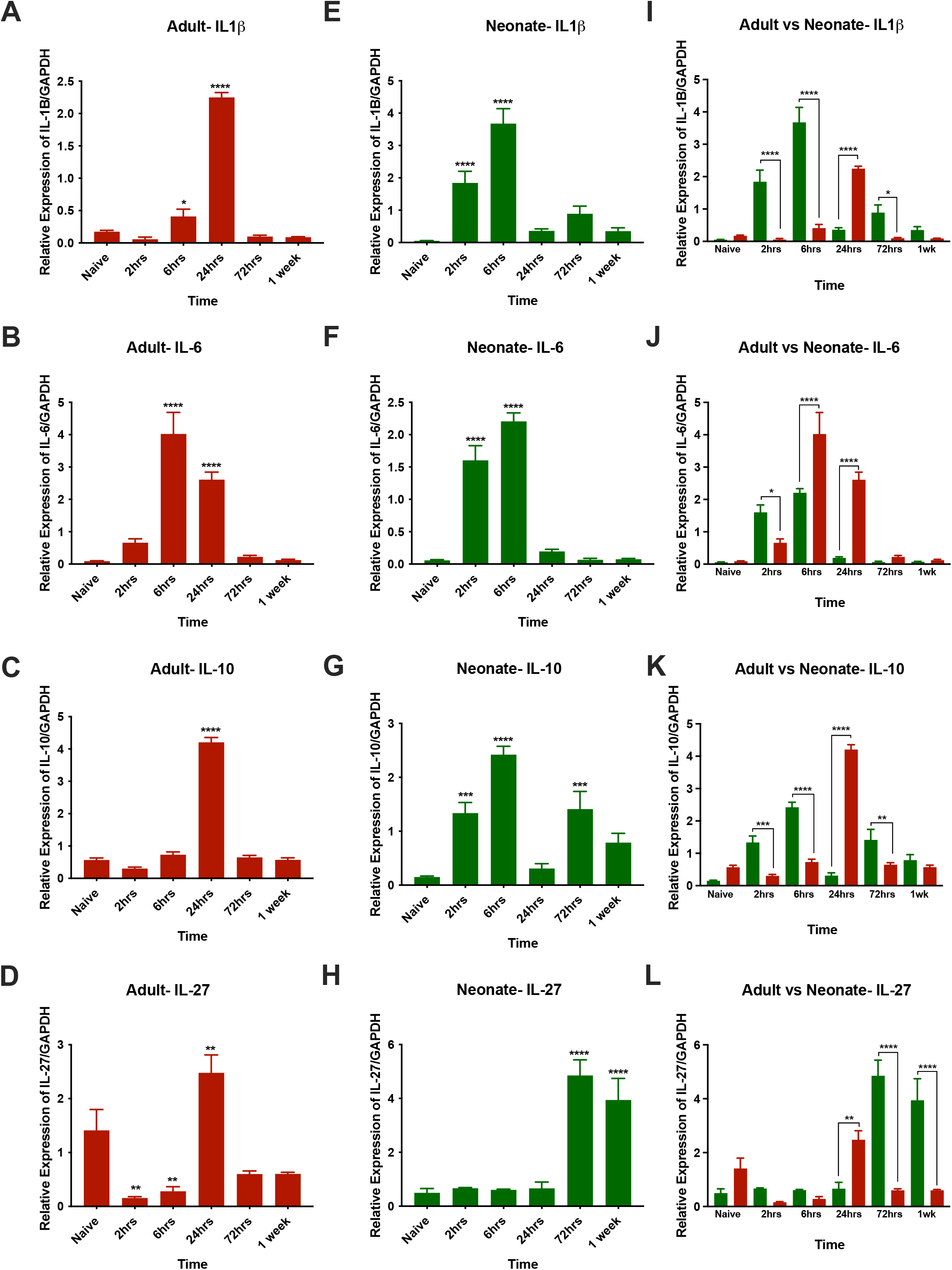
The profile of cytokine production in inflamed skin differs with age. Assessments of the expression of IL-1b (A, E & I), IL-6 (B, F & J), IL-10 (C, G & K) and IL-27 (D, H & L) following s.c. injection of CFA in neonates (green) and adults (red) were made and demonstrated differences between the age groups. Expression of all four of these cytokines were significantly increased at both ages when compared to baseline, however increases in neonates seems to precede those in adults. The magnitude of the increase seems to also differ with age with neonates producing more IL-1b and IL-27 than adults and the reverse being true for IL-6 and IL-10. Data displayed are mean ± S.E.M. Comparisons between timepoints and ages were made using a Two-way ANOVA with Bonferroni Multiple Comparison test. ****P<0.0001.

**Figure 6:**
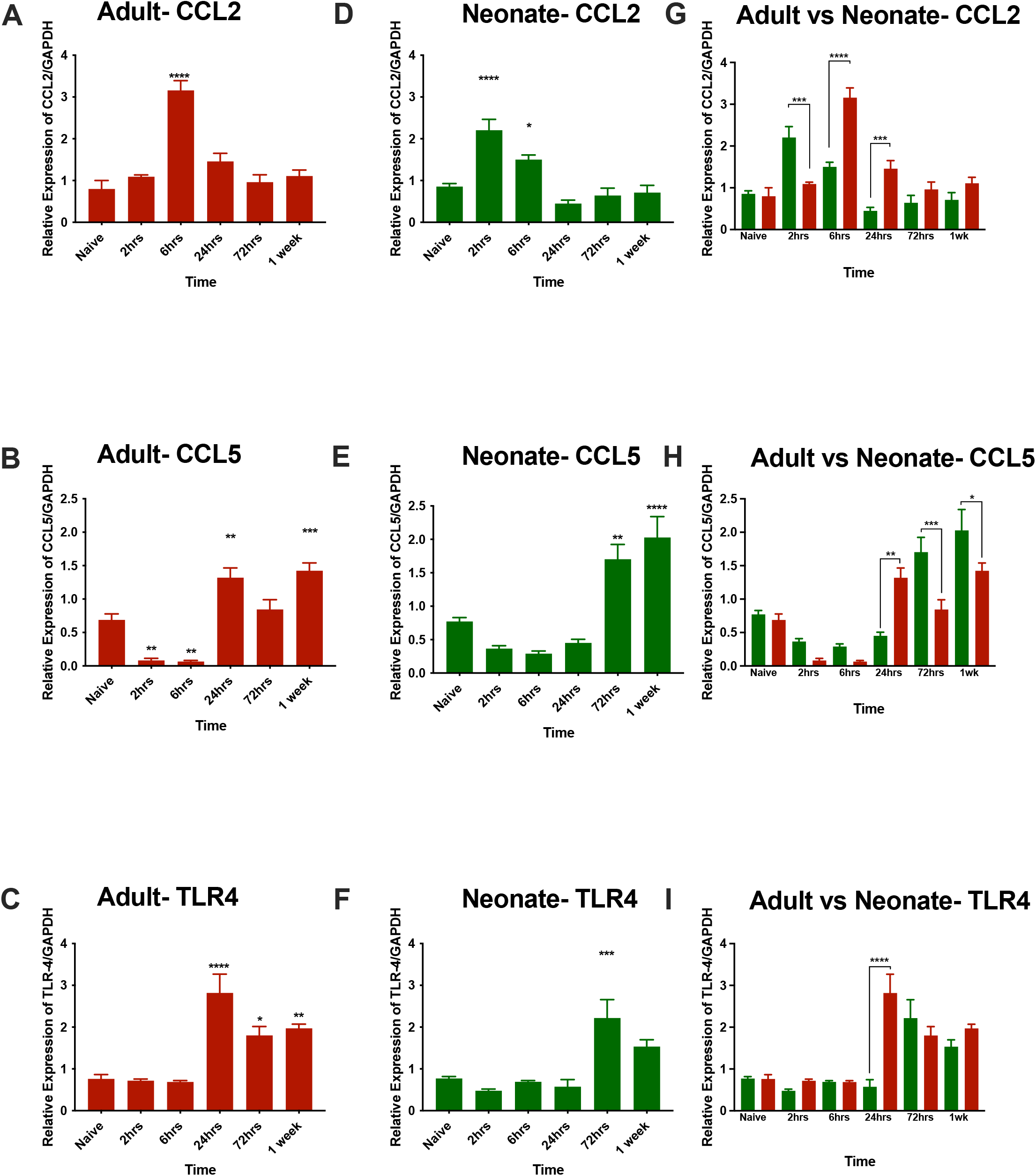
Chemokine and TLR4 expression are also regulated in an age-dependent manner. Basal levels of CCL2 (A, D & G) were comparable between the age groups as were levels of CCL5 (B, E & H) and TLR4 (C, F & I). Changes in the magnitude and time-course of changes in the expression of these were not as different as the cytokines measured. Data displayed are mean ± S.E.M. Comparisons between timepoints and ages were made using a Two-way ANOVA with Bonferroni Multiple Comparison test. ****P<0.0001.

In neonatal animals IL-1β and IL-6 mRNA expression was increased in paw tissue very rapidly following CFA injection (Fig 5e&f). Significant increases in IL-1β expression were observed 2 hours and 6 hours post-CFA injection (Control: 0.05 ± 0.01, 2 hours: 1.8 ± 0.36, 6 hours: 3.7 ± 0.46. F (5, 31) = 32.74, P<0.0001). IL-6 expression followed a similar pattern, with significant increases in expression at both 2 hours and 6 hours (Control: 0.06 ± 0.01, 2 hours: 1.6 ± 0.22, 6 hours: 2.2 ± 0.13. F (5, 27) = 86.58, P<0.0001). Expression of both cytokines quickly resolved and by 72 hours, expression of both genes returned to levels comparable to those observed in control animals. In neonates, increased expression of the anti-inflammatory cytokine IL-10 was observed at 2 hours and 6 hours post-CFA injection (Control: 0.15 ± 0.02, 2 hours: 1.34 ± 0.2, 6 hours: 2.42 ± 0.15. F (5, 29) = 20.85, P<0.0001). The IL-10 response appeared to be biphasic, as there was a significant increase in expression early on, which was followed by a reduction in expression levels at 24 hours (Fig 5g). This was followed by a significant increase in expression at 72 hours (1.41 ± 0.33 F(5,29)=20.85 P<0.001) and another reduction in expression levels at 1 week post-CFA injection. There were no significant differences in IL-27 expression levels in neonates until 72 hours and 1 week post-CFA injection in neonate animals (Naïve: 0.50 ± 0.16, 72 hours: 4.9 ± 0.58, 1 week: 3.9 ± 0.80. F (5, 31) = 22.76, P<0.0001) (Fig 5h).

Neonatal and adult animals have statistically similar levels of IL-1β in naïve animals (Neonates: 0.05 ± 0.01, Adults: 0.17 ± 0.02). However, comparisons of expression post-CFA injection showed that neonates have significantly increased IL-1β expression compared with the adults at 2 hours (Neonates: 1.8 ± 0.36, Adults: 0.06 ± 0.03) and 6 hours (Neonates: 3.7 ± 0.46, Adults: 0.4 ± 0.11). At 24 hours, adults had significantly increased expression (Neonates: 0.4 ± 0.06, Adults: 2.3 ± 0.07). At 72 hours, neonates again showed significantly increased expression (Neonates: 0.9 ± 0.24, Adults: 0.1 ± 0.02) and at 1 week post-CFA injection there were no differences in expression levels between the ages (F (5, 58) = 39.42, P<0.0001) (Fig 5I).

IL-6 expression in neonates was not significantly different at baseline but increased at 2 hours post-CFA and when compared with the adult group (Neonates: 1.6 ± 0.23, Adults: 0.7 ±.0.12). At 6 hours adults had significantly increased expression compared with neonates (Neonates: 2.2 ± 0.13, Adults: 4.0 ±.0.67). At 24 hours, expression remained significantly increased in adults and there is a huge reduction in neonate IL-6 expression (Neonates: 0.2 ± 0.03, Adults: 2.6 ±.0.23). From 72 hours onwards there were no differences in neonate and adult IL-6 expression (F (5, 56) = 15, P<0.0001) (Fig 5j).

IL-10 expression did not differ between naïve neonates and adults. At 2 hours and 6 hours, the neonate group showed significantly increased expression over the adults (2 hours – Neonates: 1.3 ± 0.2, Adults: 0.3 ± 0.04, 6 hours – Neonates: 2.4 ± 0.15, Adults: 0.7 ± 0.09). At 24 hours the adult group had significantly increased expression compared with neonates (Neonates: 0.30 ± 0.09, Adults: 4.2 ± 0.15). At 72 hours the neonates again displayed increased expression compared with adults. Neonates: 1.41 ± 0.33, Adults: 0.65 ± 0.06). At 1 week post-CFA injection there were no differences in expression levels between the two ages (F (5, 56) = 89.06, P<0.0001) (Fig 5k).

Adult rats appeared to have higher levels of IL-27 than neonates, although this difference was not significant. Expression levels at 2 and 6 hours were similar between the ages. At 24 hours, adults showed significantly increased expression over neonates (Neonates 0.7 ± 0.23, Adults: 2.5 ± 0.33). This was not maintained, as at 72 hours onwards neonates had significantly increased expression of IL-27 compared to adults (Neonates:4.9 ± 0.58, Adults: 0.6 ± 0.06) (F (5, 60) = 25.39, P<0.0001). By 1 week post-CFA injection the levels of expression of IL-27 in neonates and adults were comparable (Fig 5l).

### 3.5 Gene expression differences in chemokine and TLR expression at the site of inflammation in neonates and adults

#### 3.5.1 Chemokine expression in adults

In adults, CCL2 expression was only significantly increased at 6 hours (Naïve: 0.8 ± 0.20, 6 hours: 3.2 ± 0.23 F (5, 29) = 23.69, P<0.0001). At all other time points CCL2 expression was comparable to expression levels observed in naïve animals (Figure 7a). CCL5 expression at 2 hours and 6 hours post-CFA injection was significantly supressed (Naïve: 0.7 ± 0.09, 2 hours: 0.08 ± 0.03, 6 hours: 0.07 ± 0.02). but at 24 hours, expression was significantly increased (24 hours: 1.3 ± 0.15). At 72 hours, expression levels were reduced, but there were no significant differences between CCL5 levels in naïve animals and those animals that received CFA injection. At 1 week, CCL5 expression was again significantly increased compared with naïve animals (1 week: 1.4 ± 0.12) (F (5, 28) = 29.43, P<0.0001) (Fig 7b). In adults, expression levels of TLR-4 were significantly increased compared with naïve animals from 24 hours post-CFA injection (Naïve: 0.8 ± 0.10, 24 hours: 2.8 ± 0.45), this increase was maintained for 72 hours (1.8 ± 0.21) and 1 week post-CFA injection (2.0 ± 0.1) (F (5, 28) = 15.35, P<0.0001) (Fig 7c).

#### 3.5.2 Chemokine expression in neonates

In neonates, CCL2 was significantly increased at 2 hours and 6 hours post-CFA injection (Naïve: 0.9 ± 0.08, 2 hours: 2.2 ± 0.26, 6 hours: 1.5 ± 0.11. F (5, 29) = 17.71, P<0.0001). From 24 hours onwards CCL2 expression was reduced and levels are comparable to those seen in naïve animals (Fig 7d). Yet at this age there were no changes in CCL5 expression until 72 hours post-CFA injection, here expression was significantly increased and these levels further increased at 1 week post-CFA injection (Naïve: 0.8 ± 0.06, 72 hours: 1.7 ± 0.22, 1 week: 2.0 ± 0.31 F (5, 28) = 19.37, P<0.0001) (Fig 7e). TLR-4 expression only significantly increased at 72 hours post-CFA injection (Naïve: 0.8 ± 0.05, 72 hours: 2.2 ± 0.44. F (5, 29) = 10.52, P<0.0001). Expression levels at all other time points were not significantly different from expression levels in naïve animals (Fig 7f).

#### 3.5.3 Chemokine expression compared between neonates and adults

Comparisons between the age groups failed to identify a consistent difference. CCL2 expression levels in naïve neonates and adults were similar, with no significant differences observed. At 2 hours, neonates displayed significantly increased expression over adults (Neonates: 2.2 ± 0.26, Adults: 1.1 ± 0.04) but at 6 hours, adults had significantly increased (Neonates: 1.5 ± 0.11, Adults: 3.2 ± 0.23). At 24 hours, expression levels in adults remain significantly increased (Neonates: 0.4 ± 0.08, Adults: 1.5 ± 0.19. F (5, 58) = 15.1, P<0.0001). At 72 hours and 1 week, expression levels were similar between the age groups and no significant differences were observed (Fig 7g). Circulating levels of CCL5 in naïve neonates and adults were similar again. At 2 hours and 6 hours, there were no differences in expression levels but at 24 hours, expression of CCL5 in adults was significantly increased compared with neonatal animals (Neonates: 0.5 ± 0.05, Adults: 1.3 ± 0.15). At 72 hours neonates had significantly increased expression levels of CCL5 (Neonates: 1.7 ± 0.22, Adults: 0.8 ± 0.15) and this is maintained through 1 week (Neonates: 2.0 ± 0.31, Adults: 1.4 ± 0.12) (Fig 7h). Comparisons of TLR-4 expression show that expression levels were once again very similar between the groups, as well as 2 hours and 6 hours post-CFA injection. The only statistically significant difference between the groups occurred at 24 hours, where adult expression levels of TLR-4 were significantly increased compared with neonates (Neonates: 0.6 ± 0.17, Adults: 2.8 ± 0.45. F (5, 57) = 9.622, P<0.0001). However at 72 hours and 1 week, expression levels of TLR-4 were once again comparable between the groups (Fig 7I).

## 4. Discussion

Previous studies of responses to acute inflammatory stimuli in the neonatal period had observed a lack of behavioural hypersensitivity in the youngest animals (Walker et al., 2003), a difference from adult hyperalgesia that we confirmed in our experimental model (Figure 1). Here we show that acute immunocellular responses to inflammogens are different in the neonatal period, as are the profile of cytokines and chemokines released.

### 4.1 CFA injection caused an increased number of immune cells in neonatal paw tissue and increased expression of CD68^+^ MR^+^ cells

During inflammation, macrophages exhibit a predominantly pro-inflammatory (M1) phenotype or an anti-inflammatory (M2) phenotype depending on the environment. M1 macrophages promote T helper (Th) 1 responses, which are pro-inflammatory. CD4^+^ T cells release interferon (IFN)- and interleukin (IL)-2, that in turn activate B cells to trigger an adaptive immune response. M2 macrophages promote tissue repair and Th2 responses. These cells are responsible for production of IL-4, IL-6, IL-10 and IL-13 (Berger, 2000). Given that it is widely accepted that neonatal immune responses are skewed to an anti-inflammatory phenotype (Chen and Field, 1995; Forsthuber et al., 1996), we hypothesised that there would be differences in the profile of macrophages recruited during inflammatory insult, which we observed.

In the naïve state there were significantly more resident immune cells present in neonatal paw skin (21.3 cells per field of views) as compared with adult paw skin (16.6 cells per field of view). Given that the neonatal immune system is considered somewhat immature (Georgountzou and Papadopoulos, 2017; Simon et al., 2015), it might seem intuitive that there would be fewer tissue-resident immune cells, but this was not the case. The naïve adaptive immune system of neonates, means they rely heavily on innate immune cells as their first-line of defence against pathogens (Maródi, 2006). The distinct bias towards a Th2 environment and lack of Th1 cytokine productions leaves them vulnerable to developing adverse reactions to microorganisms extremely quickly (Cuenca et al., 2013). The higher number of tissue resident macrophages in neonatal tissue may reflect the greater need for an innate immune response in early life, to compensate for the relatively immature adaptive immune system..

Immunohistochemistry of the paw skin showed that in adults there was significantly higher expression of MR positive cells at all time points, whereas in neonate animals there is significantly higher co-expression of CD68 and MR. MR is expressed on macrophages that have been alternatively activated and so possess anti-inflammatory properties. IL-4 and IL-13 upregulate MR expression on macrophages, causing endocytosis and antigen presentation (Gordon, 2003). It is interesting that MR only positive macrophages are significantly reduced in neonatal animals, which suggests under these conditions they may possess a more pro-inflammatory Th1-like phenotype. This does not suggest this is the neonatal response in all situations, indeed during neonatal sepsis the neonatal immune response is highly anti-inflammatory, mediated by IL-10 and TGF-β to limit inflammation and promote resolution (Bone et al., 1997). Other research also suggests that neonatal immune cells produce less IFN-γ, IFN-α, and IL-12 than adult immune cells (Kraft et al., 2013), all of which are key pro-inflammatory cytokines, but produce greater amounts of IL-6, IL-1β and IL-10 than adults (Kollmann et al., 2009), and may contribute to the lack of hyperalgesia in neonates. Also noteworthy, is that neonatal macrophages co-express CD68 and MR at very high levels, significantly more so than adult macrophages. Macrophages that express both these markers are still considered M2 macrophages and therefore anti-inflammatory (Boytard et al., 2013).

### 4.2 Altered macrophage profiles are not as a result of altered neutrophil recruitment to the site of inflammation

As previously discussed, overall neonates had higher total numbers of immune cells present at the site of inflammation. This is in line with reports in the literature that granulocytes are the most abundant cell type in neonates (Urlichs and Speer, 2004) and are circulating in greater numbers during the first 24 hours after birth. However, some reports suggest that the number of circulating neutrophils in neonates is reduced. In neonatal humans, neutrophil numbers range from 1.5–28 × 10^9^ cells/L blood, whereas in adults levels of 4.4 × 10^9^/L blood are observed (Melvan et al., 2010). Here no differences in the number of neutrophils in naïve neonatal or adult rats were observed; neutrophil numbers are significantly increased 2 hours post-injection in both age groups and thus are not likely to be responsible for the lack of behavioural hypersensitivity seen in younger animals.

### 4.3 Neonates have increased IL-10 expression compared with adults in the first 6 hours post-CFA injection

Our data showed that IL-10 was expressed at significantly higher levels in neonates compared to adults at 2 hours and 6 hours post-CFA injection. This could contribute to the prevention of onset of hyperalgesia in neonatal animals. In a nerve injury model, IL-4 and IL-10 have been shown to prevent the onset of pain in early life (McKelvey et al., 2015) however this study was conducted in the CNS. The elevated levels of IL-10 in neonates were not sustained and reduced to levels comparable with naïve animals by 24 hours post-CFA injection. Although there was clearly an initial boost in anti-inflammatory cytokine production in neonates, this did not prevent the production of pro-inflammatory cytokines. The levels of pro-inflammatory cytokines IL-1β and IL-6 were raised significantly in neonates compared to their naïve counterparts. Studies have reported that the increased production of pro-inflammatory IL-1β and IL-6 in neonates may have limited effects due to the reduced production of TNF-α and increased production of IL-10 (Kollmann et al., 2009). The same study also reported that the increased production of IL-10 by neonatal cells may have been because of the increased production of IL-6 and IL-23 (Kollmann et al., 2009). The data presented here is in agreement with this work, the relative levels of expression of IL-10 and IL-6 at 2 hours and 6 hours post CFA injection in neonatal animals are extremely similar. In summary, this early increased production of IL-10 could be key in preventing the onset of hyperalgesia in neonatal animals, however, further investigation is required.

### 4.4 Neonates and adults display different expression of CCL2 and CCL5 after CFA injection

CCL2 has been shown to drive macrophage polarisation towards an anti-inflammatory, M2 phenotype (Sierra-Filardi et al., 2014). CCL2 is also referred to as monocyte chemotactic protein-1 (MCP-1) and as its name suggests monocytes and macrophages infiltrate inflamed tissues under its influence. Once recruited, macrophages produce high levels of CCL2 (Yoshimura et al., 1989), in turn recruiting more macrophages and other immune cells including mast cells and basophils. We show that differences in CCL2 expression in neonates and adults is from 2 – 24 hours post CFA injection. Initially neonatal expression of CCL2 is significantly increased compared to adults, however, this changes at 6 hours with adults producing more CCL2 which is sustained until the 24 hour timepoint. There is little research into chemokine expression in neonates, both in rodents and humans. A study by (Winterberg et al., 2015) carried out to identify phenotypic features of neonatal macrophages in mice showed differences in CCL2 expression, amongst other cytokine and chemokines.

As with CCL2, there is little available research into the differences between neonatal and adult expression of CCL5. A study carried out in 2012, showed that preterm human neonates produced significantly lower levels of CCL5 than term neonates, however this data was not compared to adult levels of CCL5 (Królak-Olejnik and Olejnik, 2012). CCL5 is important in recruitment of other immune cells to the site of inflammation, including macrophages, eosinophils, basophils and T cells. IL-2 and IFN-γ are released by T cells, these interact with CCL5 and induce production of natural killer (NK) cells, which in turn generates C-C chemokine-activated killer cells (Soria and Ben-Baruch, 2008).

### 4.5 Conclusion

In conclusion, changes in the cellular morphology taken together with the more rapid production of IL-1β, IL-6 and IL-10 expression in neonates suggests that neonatal animals have a more rapid response to CFA injection that is skewed towards an M2 phenotype. This may begin to provide an understanding as to why hyperalgesia is not observed in neonates following CFA injection.

